# Retina Gap Junction Networks Facilitate Blind Denoising in the Visual Hierarchy

**DOI:** 10.1101/2022.05.16.491952

**Authors:** Yang Yue, Kehuan Lun, Liuyuan He, Gan He, Shenjian Zhang, Lei Ma, Jian.K. Liu, Yonghong Tian, Kai Du, Tiejun Huang

## Abstract

Gap junctions in the retina are electrical synapses, which strength is regulated by ambient light conditions. Such tunable synapses are crucial for the denoising function of the early visual system. However, it is unclear that how the plastic gap junction network processes unknown noise, specifically how this process works synergistically with the brain’s higher visual centers. Inspired by the electrically coupled photoreceptors, we develop a computational model of the gap junction filter (G-filter). We show that G-filter is an effective blind denoiser that converts different noise distributions into a similar form. Next, since deep convolutional neural networks (DCNNs) functionally reflect some intrinsic features of the visual cortex, we combine G-filter with DCNNs as retina and ventral visual pathways to investigate the relationship between retinal denoising processing and the brain’s high-level functions. In the image denoising and reconstruction task, G-filter dramatically improve the classic deep denoising convolutional neural network (DnCNN)’s ability to process blind noise. Further, we find that the gap junction strength of the G-filter modulates the receptive field of DnCNN’s output neurons by the Integrated Gradients method. At last, in the image classification task, G-filter strengthens the defense of state-of-the-arts DCNNs (ResNet50, VGG19 and InceptionV3) against blind noise attacks, far exceeding human performance when noise is large. Our results indicate G-filter significantly enhance DCNNs’ ability on various blind denoising tasks, implying an essential role for retina gap junction networks in high-level visual processing.

## Introduction

A fundamental challenge for the early visual system is noise reduction. In the outermost layer of our visual system, photoreceptors (rods) detect light signals entering the retina with single-photon resolution ***Rieke and Baylor (1998***). However, this process suffers from considerable noise interference originating from ambient environment light resources and multiple intrinsic biological processes ***Barlow et al. (1993); Angueyra and Rieke (2013***). The situation of mixed noise origination faced by the retina is equivalent to the long-standing issue in the machine learning field called “blind noise reduction”, which aims to remove noises of unknown type ***Yue et al. (2019***). Various advanced methods have been developed in both academia and industry to achieve more general noise reduction ***Dabov et al. (2006); Chen et al. (2018); Khmag et al. (2018); Yue et al. (2019***). However, their performances are still dependent on a broader prior estimation of noise type and intensity.

Compared to machine learning, the biologically evolved image denoising strategy discovered in the retina provides an inspiring solution to this problem. Over millions of years of evolution, the retina has developed a variety of cellular and subcellular denoising mechanisms ***Demb et al. (2004); DeVries et al. (2002); Van Rossum and Smith (1998***). Among all these mechanisms, gap-junction coupling has been the major denoising strategy to tackle unknown noise, and no prior estimation of noise type or intensity seems to be required ***Smith and Vardi (1995***). Contrary to chemical synapses, gap junctions are electrical synapses, allowing bi-directional and unattenuated currents to pass through between adjacent cells ***Bloomfield and Völgyi (2009); Evans and Martin (2002); Li et al. (2012***). This property makes gap junctions capable of achieving rapid electrical signal synchronization around the local area in a network, thus contributing to smoothing the signals ***DeVries et al. (2002); Li et al. (2012***). Interestingly, the retina gap junction also exhibits unique plasticity, when ambient illumination conditions alter. Specifically, because the impact of noise becomes more dramatic in dim light due to the decrease in signal-to-noise ratio, gap junction conductance can vary accordingly within the day and night cycles, generating adaptive denoising strength ***Bloomfield and Völgyi (2009); Xin and Bloomfield (1999); Yao et al. (2018***).

Another conceptual gap is how retina denoising strategies impact higher visual centers. It has become clear that the information processing in the visual system is hierarchical ***Brocas and Carrillo (2008); Meunier et al. (2009***). Light signals first transcode in the retina, then relay to lateral geniculate nuclei (LGN), and finally reach the visual cortex, which spans low-level information processing and high-level cognition, termed visual hierarchy ***Lennie (1998); Lu et al. (2018***). Due to the limitation of experimental methods, it is infeasible to study the retina and cortex in the visual hierarchy simultaneously. Recent fMRI studies show that the early visual areas (V1 and V2) may have a functional role related to low-level 2D visual tasks such as denoising ***Pratte et al. (2013); Dwivedi et al. (2021***), whereas later visual areas (V3, V4, and IT) may involve 3D and semantic tasks such as classification ***Eickenberg et al. (2017***). Although those experiments indicate the associations between the retina and high order visual modules in denoising, the computational mechanism of the earlier visual cortex in denoising is not fully explored. In particular, how such cortical mechanisms are in synergy with retina denoising strategies in the visual hierarchy remains unclear.

To computationally investigate the impact of denoising effects of the retina on higher modules in the visual hierarchy, we utilize deep convolutional neural networks (DCNNs) as a model of the primate ventral visual pathway. The contemporary architectures of DCNNs are closely related to the visual hierarchy, reflecting the workflow of the primate ventral visual pathways ***Güçlü and van Gerven (2015***). Hence, DCNNs as brain models in cognitive neuroscience are widely used in simulating visual denoising ***Pratte et al. (2013***), emotion perception ***Zhou et al. (2021***), face and object classification ***Cadieu et al. (2014); Eickenberg et al. (2017***). Further Researches on cognitive neuroscience show that DCNNs have a high functional similarity with primate ventral visual pathways ***Cichy et al. (2016); Eickenberg et al. (2017); Seibert et al. (2016***). Therefore, we can leverage DCNNs as computational models of the primate ventral visual pathways to quantitatively study the influence of the lower layer’s denoising effect on visual cognitive tasks performed by higher layers in the visual hierarchy.

In this work, inspired by our observations from electrically coupled photoreceptors, we construct a gap-junction filter (G-filter) for blind noise reduction. We quantitatively assess the blind denoising performance of the G-filter on various types of noise and find that G-filter is a general-purpose blind denoiser, transforming the noise distribution into a similar form. To further explore the influences of the gap junction network in the retina on high-level visual functions, we adopt denoising convolutional neural network (DnCNN) ***Zhang et al. (2017***) as the primary visual cortex and combine it with G-filter, called Hybrid DnCNN. Intriguingly, G-filter dramatically elevates the DnCNN’s ability in blind denoising. Furthermore, we show by the integrated gradients method ***Sundararajan et al. (2017***) that the neurons’ receptive fields in the Hybrid DnCNN are influenced by the gap junction conductance of the G-filter, which implies that the deep learning model automatically adapts to the plasticity of G-filter. Finally, based on a recent psychophysical experiment of image classification task ***Geirhos et al. (2018***), We show that G-filter significantly improves the ability of state-of-the-art DCNNs to resist blind noise attacks, even far outperforming humans under big blind noise.

In summary, our work demonstrates that the gap junction network is essential for image denoising in the retina and enhances higher visual functions, object recognition, and learning because it provides the denoised information to the higher hierarchy. Notably, the gap junction networks we developed can be directly merged into the modern AI architecture, bringing more convenience for machine learning to integrate biologically-inspired blind denoising.

## Results

### Plastic retina gap-junction network in the visual hierarchy

Light signals from the natural environment always carry tremendous noise after entering the retina ***Schneeweis and Schnapf (1999); Passaglia and Troy (2004***). However, the visual representation in our minds is fairly fluid and clear. Light signals must undergo intense noise reduction as they traverse the entire visual pathway, whether in the outmost layer of the visual system — the photoreceptor layer or the higher visual cortex ***Ruda et al. (2020***) (Fig. 1A). To investigate the denoising effect of the first layer of the signal relay, we establish a biologically detailed photoreceptor network model (see “Methods”). In the retina, neighboring photoreceptors are directly connected by gap junctions, enabling information sharing between neighbors (Fig. 1B, C, see “Methods”). As a consequence, each photoreceptor may not only respond to the light stimulus captured by its own cell, but also to the light stimulus received by adjacent cells, which can also be seen as a receptive field. The commonly used method to compute the receptive field is spike-triggered average (STA) ***Chichilnisky (2001***), also known as “white-noise analysis”, which has been widely applied to explore receptive fields of neurons in the visual pathway***Chichilnisky (2001); Rust et al. (2004); Casti et al. (2008***). Here, we use STA to generate neurons’ receptive field under light stimulations in the photoreceptor network.

**Figure 1.**
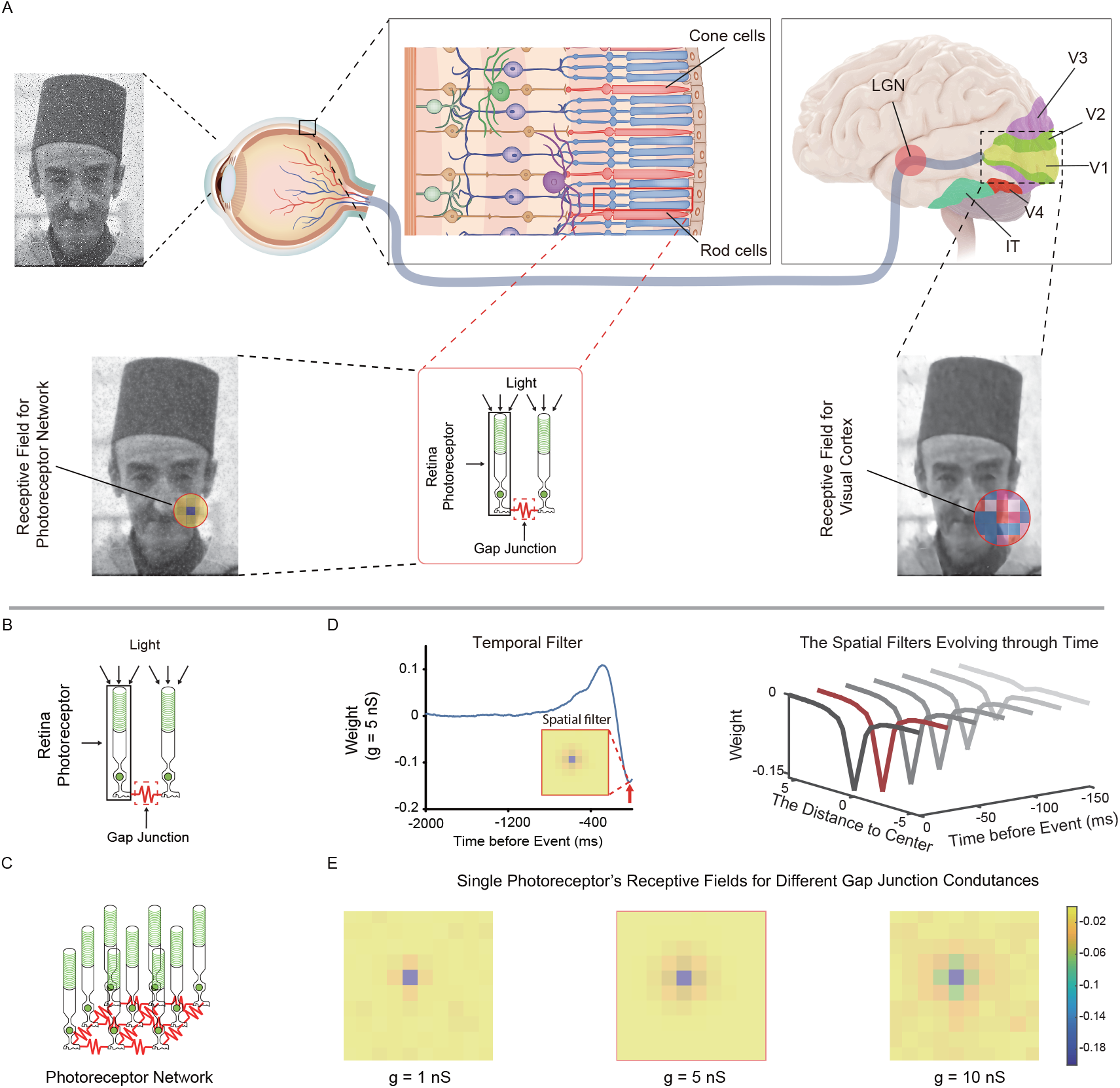
Plastic receptive fields in the visual hierarchy. (A) Information processing in the human visual system is hierarchical (retina - LGN - cortex), where the image reconstruction and recognition in the cortex may be related to the ventral visual pathway (from V1 to IT). As the layers deepen, the receptive fields of neurons become larger and more complex. (B) Photoreceptors are connected by a gap junction, where the photoreceptors are located at the first layer of the visual hierarchy. (C) The photoreceptor network with grid-like gap junctions. (D) The spatiotemporal filter (receptive field) of the 10 × 10 photoreceptor network evaluated by STA. (E) Plastic receptive fields in the 10 × 10 photoreceptor network, where the gap junction conductances are 1 nS, 5 nS, and 10 nS, respectively. The portrait in our figures comes from a public dataset - BSD68 Martin et al. (2001).

To compute the STA in a 10 × 10 photoreceptor network with gap junctions, we set the white noise to 5Hz as network input and record the responses (see “method”). The STA of a single photoreceptor in the network exhibits the spatiotemporal filtering effect (‘spatiotemporal receptive field’). For example, at the temporal level, the network shows moderate inhibition immediately before an event occurs (red arrows in Fig. 1D), which is attributed to negative photocurrents evoked by the stimulus and gap junction connections in the network. Over time, the inhibitory effect of the spatial filter gradually increases at each time step (Fig. 1D, right).

As in the retina, the connection strengths of neighboring photoreceptors are affected by the change of the gap junctions’ conductance ***Bloomfield and Völgyi (2009***), we here adjust the conductance to see its effect on spatial receptive field. The values of gap junction conductance (1*nS*, 5*nS* and 10*nS*) are within the experimentally reported range ***Rozental et al. (1998); Zhang and Wu (2005); Jacoby et al. (2018***). By increasing the gap junction conductance from 1*nS* to 10*nS*, the receptive field range of the photoreceptor cell is slightly enlarged, while the weights between adjacent cells are significantly enhanced, indicating an increased contribution of signals from similar neighbors (Fig. 1E). Integrating more information from neighbors may cause the signals to be locally smoothed, providing the earliest noise reduction procedure. Furthermore, with larger receptive fields, individual photoreceptors may contain more precise statistics about noise distribution and signal patterns ***Balasubramanian and Sterling (2009***), helping to remove noise and preserve useful information for the higher visual hierarchy.

### Gap junction filter (G-filter) is a general-purpose denoiser

To apply the photoreceptor network to machine learning architectures, we develop a gap junction filter (G-filter) and make some modifications. Specifically, we keep the gap junction connections in the previous network but alter the detailed photoreceptor model in the following ways. We remove all ion channels in photoreceptors except the passive leak channel. The single neuron model in the gap junction filter is described as:

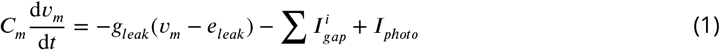

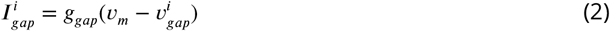

where *C*_*m*_ is the membrane capacitance, *v*_*m*_ is the membrane voltage, *g*_*leak*_ is the conductance of passive leak channel, *e*_*leak*_ is the potential of passive leak channel, 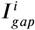 is the *i* − *th* gap junction current, 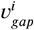 is the *i* − *th* neighboring membrane voltage and *I*_*photo*_ is the photocurrent. We use a double-exponential function as photocurrent to replace the phototransduction cascade model.

The photocurrent is defined as:

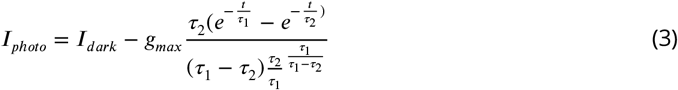

where *I*_*a k*_ = 40*p, τ*_1_ = 64*m, τ*_2_ = 68*m*, and *g*_*ma*_ controls the maximum amplitude of *I*_*photo*_.

The gap junction filter structure is still grid-like as in the previous model (Fig. 1C). Here, we let the size of the network expands according to image resolutions, wherein each photoreceptor encodes a single pixel of the input image (Fig. 2A). At first, each pixel as a photo signal is transformed into a current curve by Eq.3, where *g*_*max*_ equals the value of the pixel. Then, the current is injected into the neuron, and the voltage curve of the neuron is recorded. The negative peak value of the voltage curve is denoted as the neuron’s output. Finally, all output values are normalized into [0, 1] as the denoised image (see “Methods”).

**Figure 2.**
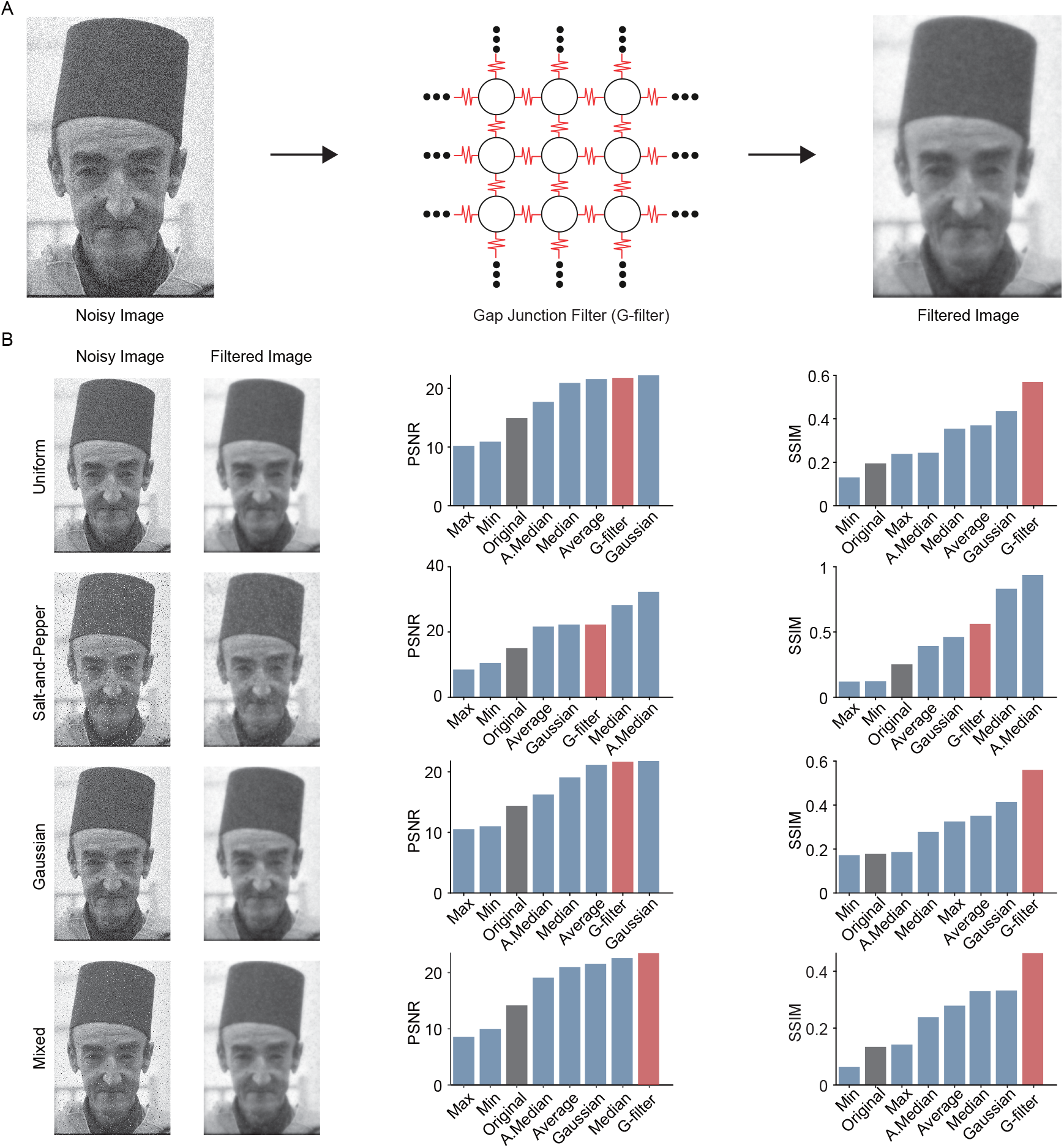
Gap junction filter (G-filter) is a general-purpose denoiser. (A) The architecture and workflow of the G-filter. Each pixel in the image corresponds to an individual photoreceptor, and the noise is filtered by gap junctions. (B) Performance of different denoising methods. On the left are the noisy samples (Uniform Salt-and-Pepper, Gaussian, and Mixed noise) processed by the G-filter. In the middle is the denoising performance of the G-filter compared to the classic noise filter under the PSNR. On the right is the denoising performance at the SSIM index. The black, blue, and red bars represent the original image qualities, the image qualities processed by the classic noise filter, and the image qualities processed by the G-filter, respectively.

To evaluate the performance in blind denoising of the G-filter, we first test it on the Berkeley segmentation dataset (BSD68) ***Martin et al. (2001***) and compare it with simple filters, including adaptive median, average, Gaussian, max, median, and min filters (see “Methods”). We apply these filters to multiple regular noise types, such as Gaussian, Laplacian, Salt-and-Pepper, and Uniform noise, plus an artificially designed mixture of noise as “blind noise”. This mixture of noise mingles all noise types mentioned above, where the intensities of each type are specified, and the proportions are random, resembling the noises with various sources generated in the natural visual processing. The original and filtered images comparison is shown in Fig. 2B (left). Since our purpose is to assess the generality of all these filters and not to pursue optimal values, we set the parameters of all simple filters to MATLAB’s default values (see Methods). Likewise, we set the gap junction conductance of the G-filter to 5 nS, which is the median value within our set range. When the filter parameters are fixed, the stability performance of the filer at different noise types and intensities reflects its generality.

To measure the denoising effect of all these noise filters, we adopt two well-known indices of image quality ***Hore and Ziou (2010***), i.e., peak signal-to-noise ratio (PSNR) and structural similarity index measure (SSIM) (see Methods). PSNR is a commonly used method to evaluate the image reconstruction quality, which computes the ratio between the maximum power of a signal and the power of noise ***Hore and Ziou (2010***). By contrast, SSIM is a perception-based method for predicting the perceived quality of an image ***Wang et al. (2004); Hore and Ziou (2010***). In the results of the PSNR index, the performance of the G-filter consistently ranks among the top three for each noise type (Fig. 2B middle, Appendix Table1 for others). Other simple filters all have weak performances in at least one noise type. For example, median/adaptive median filters perform best on salt and pepper noise, but not well on other types of noise.

Intriguingly, on the SSIM index, we find that the performance of G-filter is dominant in nearly all noise types (Fig. 2B right; Appendix Table2 for others). Then we decided to further explore the G-filter mechanism under the SSIM index. As the total SSIM index value represents the product of luminance, contrast, and similarity (Fig. 3), we analyze the G-filter performance on each component. Unlike PSNR, SSIM allows us to compute and visualize the SSIM index at the pixel level. Analyzing the SSIM performance in an example figure processed by the G-filter, we find that the G-filter scores higher on the luminance and similarity component but relatively lower on the contrast component, indicating it is primarily used for smoothing the background and the object body, but sharp textures such as outlines are blurred (Fig. 3B). This particular feature might be helpful for denoising in dim light conditions where most object detail is lost (see Discussion for more details). Overall, the G-filter is a general-purpose denoiser that is efficient for multiple noises, including the blind noise, but slightly compromising contour information.

**Figure 3.**
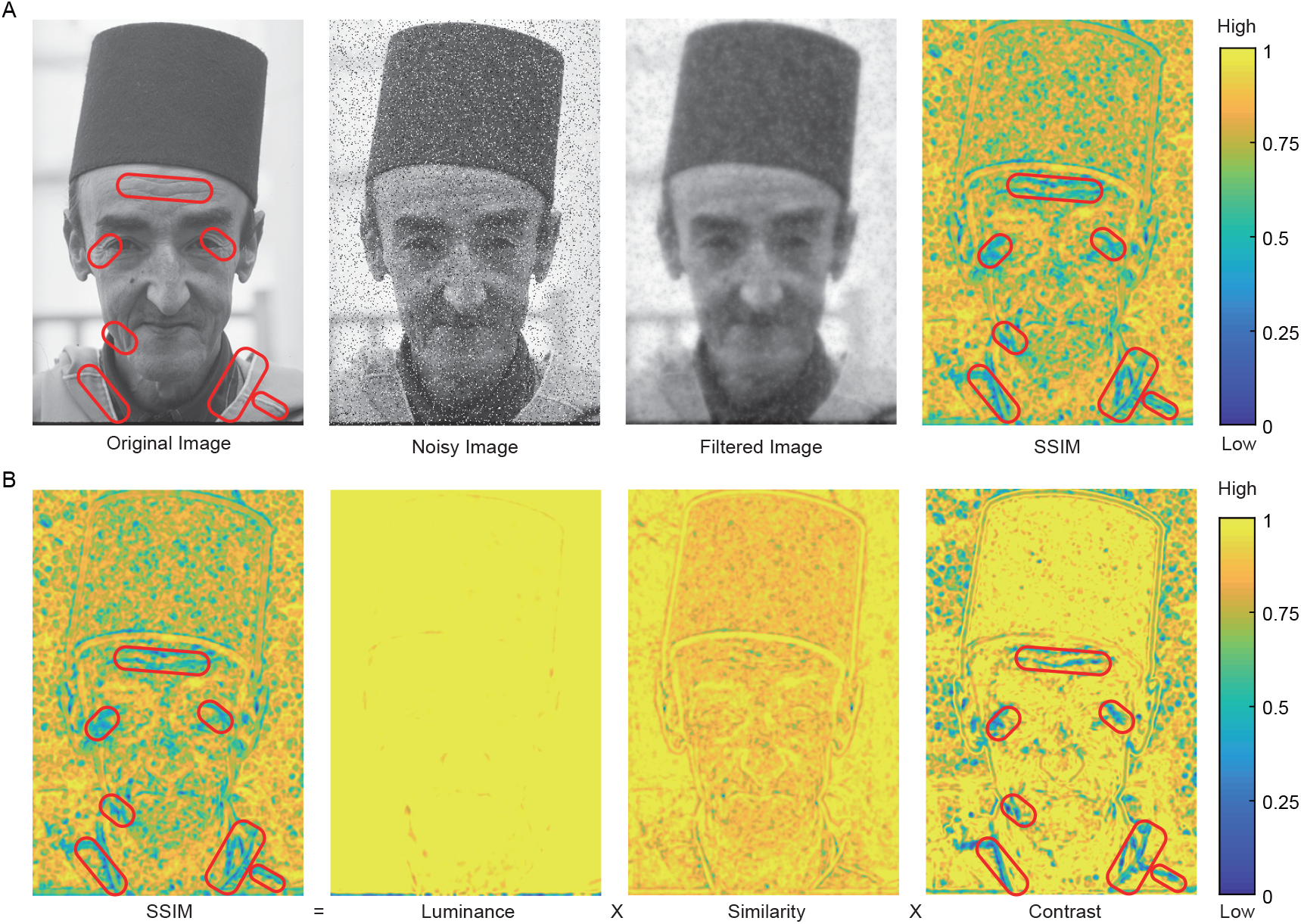
High contrast leads to the low image quality of the G-filter processed image under the SSIM index. (A) A noisy sample (Salt-and-Pepper noise) processed by the G-filter and evaluated by the pixel-wise SSIM index. The red circles represent low image quality areas. (B) The decomposition of the pixel-wise SSIM index. The SSIM index consists of luminance, contrast, and similarity, where the contrast is mainly responsible for the low image quality.

### G-filter transforms distinct noise distributions into a similar form

To further understand the mechanism of the G-filter acted upon all these types of noises, we wonder if G-filter changes the noise distributions. To test this hypothesis, we add uniform noise, Gaussian noise, Salt-and-Pepper noise, and mixed noise to one sample image and compare the noise distribution before and after G-filter application (Fig. 4A). Interestingly, we find the G-filter can change different noise distributions into a similar bell-shaped form, no matter how distinct their original distributions are. For example, the uniform noise distribution looks like a rectangle, and the Salt-and-Pepper noise distribution is at both ends of the x-axis, but their noise distribution after G-filtering is very similar in the sample image (Fig. 4C). To confirm this conclusion holds for the entire dataset, we average the noise distribution of the original and G-filtered images for the whole testing set (Fig. 4B). The results are consistent with the observations on a single image (Fig. 4D).

**Figure 4.**
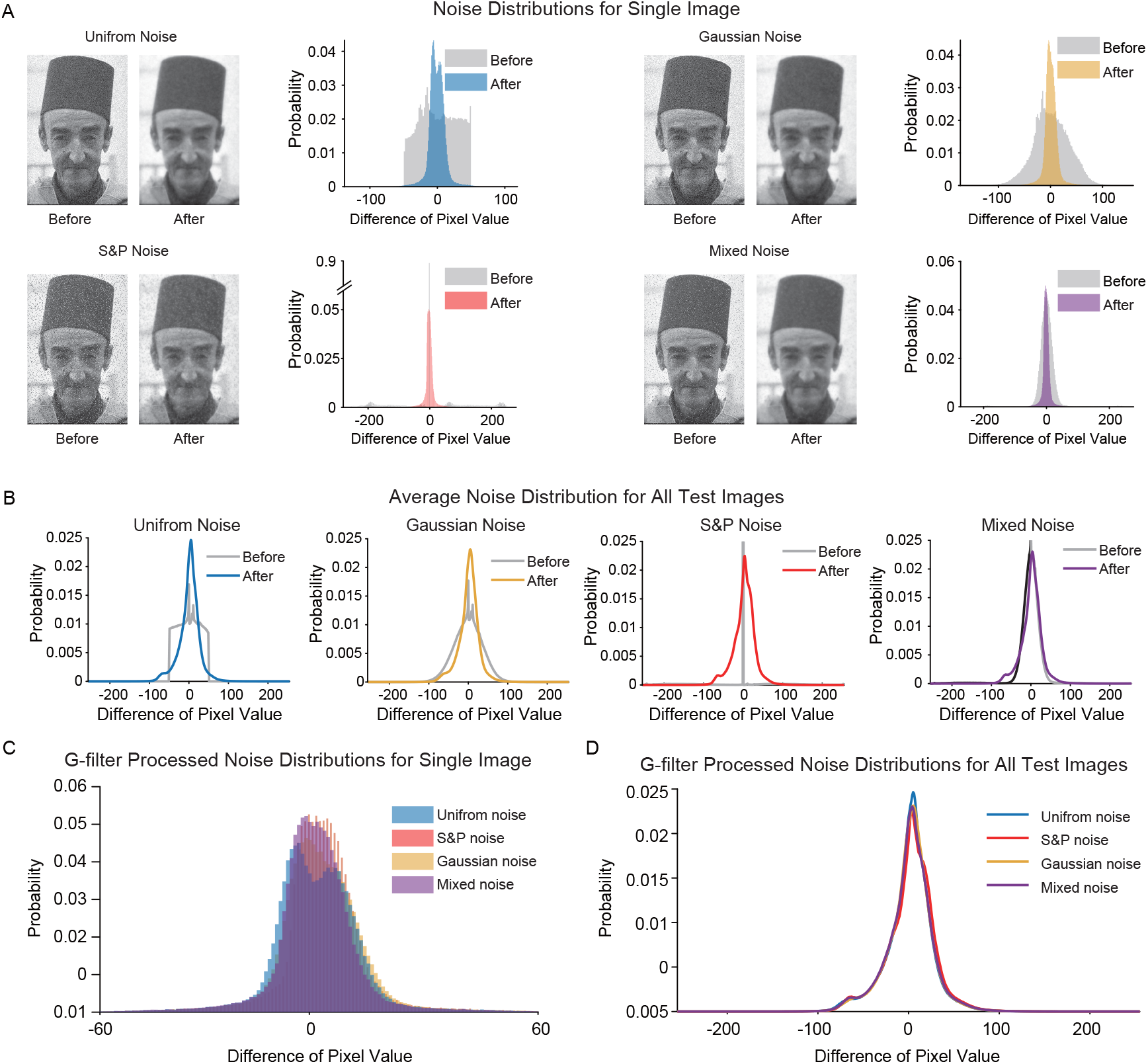
The G-filter transforms different noise distributions into a similar form. (A) Noise distributions of a single image with different noises (Uniform Salt-and-Pepper, Gaussian, and Mixed noise) before and after G-filter processing. The gray regions represent original noise distributions, and the colored regions represent G-filter processed noise distributions. (B) Noise distribution statistics of images under different noises before and after G-filtering, where the statistical range includes all test images in BSD68. Similarly, the gray curves represent the average of original noise distributions, and the colored curves represent the average of G-filter processed noise distributions. (C) The sample images with different noise types have similar noise distributions after processing by G-filter. (D) The average noise distributions of all test images with different noise types have similar noise distributions after processing by G-filter.

So far, we find that G-filter is a dynamic inhibitory filter, which can be applied to a broad range of noise types, including blind noise. Furthermore, by changing noise distributions of different types into bell-shaped form, G-filter can convert blind noise into a homogeneous pattern and potentially make it more tractable for subsequent deep learning modules.

### G-filter strengthens DnCNNs’ defense against blind noise in image denoising tasks

In previous results, we find that although G-filter is a versatile filter to various noise types, it also sacrifices some critical information such as the contours and textures. If this is also true for the retina, is it possible that our cortex can learn to compensate for the lost details? One clue is that the local loss of visual information caused by retina macular degeneration can be to an extent compensated by the visual cortex ***Baker et al. (2005); Dilks et al. (2014***) (also see Discussion for more details). Hence, we reason that the blurry contour information caused by our G -filter can also be repaired by a subsequent cortical-like deep learning architecture while preserving the noise filtering effect. To this end, we need to choose a DCNN framework as the cortical model for denoising and image reconstruction tasks. Specifically, the early visual areas (V1 and V2) are believed to have a noise-filtering mechanism ***Pratte et al. (2013***). Therefore, we choose a classic DCNN-based denoising architecture named denoising convolutional neural network (DnCNN) ***Zhang et al. (2017***) as a biologically plausible model for the early visual cortex, which resembles V1 and V2 structurally and functionally.

DnCNN is a full-convolutional and residual network consisting of several convolutional layers of the same size as input and is initially designed for image noise reduction ***Zhang et al. (2017***). The output layer of DnCNN has a skip connection, which subtracts the input image and noise as the denoised image. We combine the G-filter before the DnCNN architecture; thus, the noisy image can be processed by G-filter first and then used to train the DnCNN (Fig. 5A). This new architecture is similar to our visual system: noisy signals are first processed in the retina and then sent to the visual cortex for further interpretations. The images in the training set are added with Gaussian noise (18dB), and the images in the testing set are with Salt-and-Pepper noise (14 dB, 16 dB, 18 dB, and 20dB). The training process follows the settings in ***Zhang et al. (2017***) (see Method). Therefore, the combined model is developed as a blind denoising model to deal with situations where no prior knowledge of noise type is available, as the noise types we used in training and testing sets are different. The performance of this model can represent the blind denoising ability.

**Figure 5.**
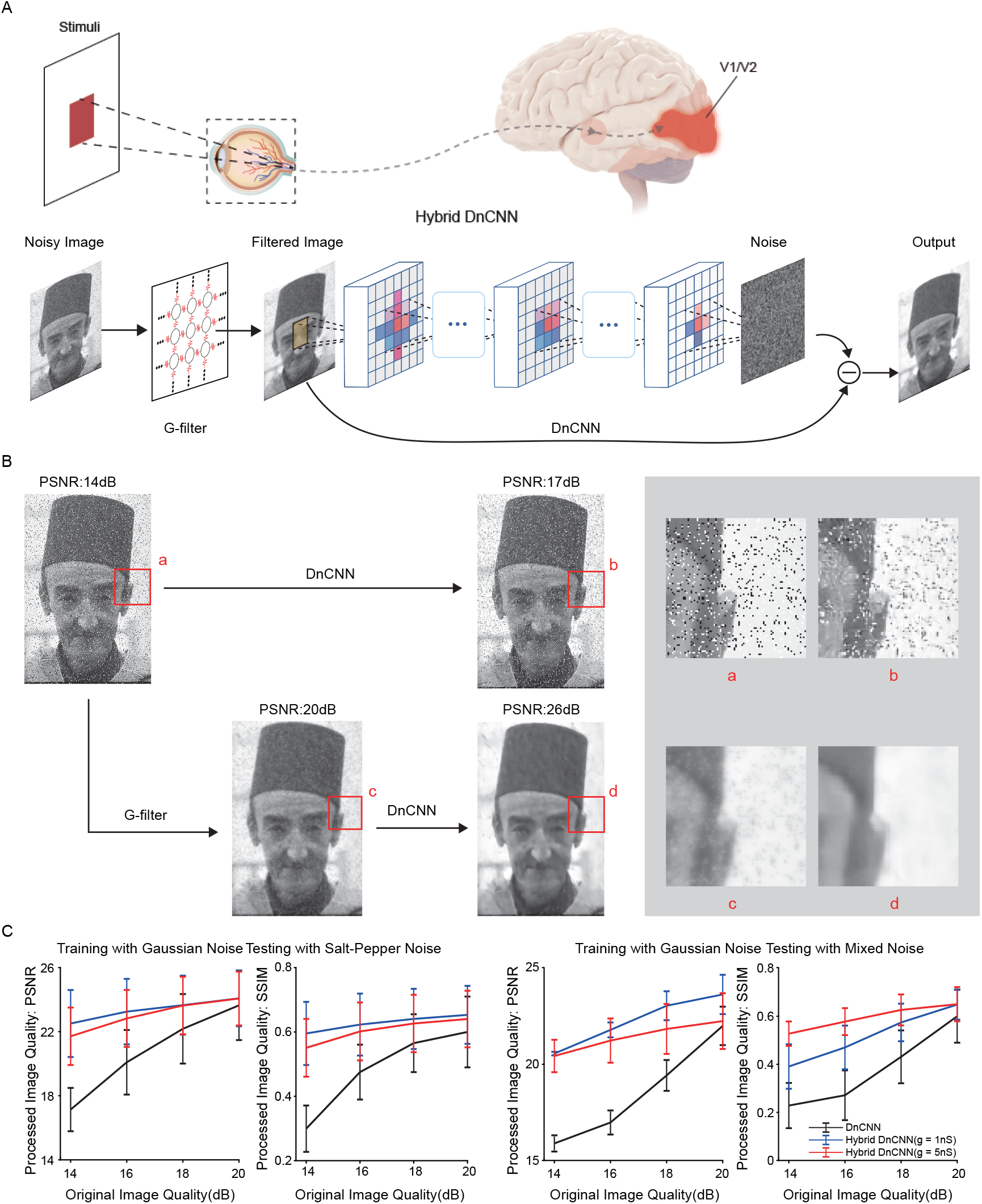
The combination of G-filter and DnCNN (Hybrid DnCNN) simulates image reconstruction (denoising) in the visual hierarchy and improves blind denoising performance. (A) The architecture and workflow of Hybrid DnCNN. The G-filter is a computational module placed in front of DnCNN for blind denoising. The G-filter is an abstraction of the retina, while the DnCNN mainly maps to V1 and V2 of the visual cortex. (B) The denoising performance of DnCNN and Hybrid DnCNN on a noisy sample. On the right is an enlarged view of the red square. (C) The denoising performance (PSNR and SSIM) of DnCNN and Hybrid DnCNN (g = 1 nS and g = 5 nS). These models are trained using a training set with Gaussian noise. On the left is the denoising performance on the testing set with salt-and-pepper noise, and on the right is mixed noise. The black, blue and red lines represent the denoised image qualities of DnCNN, Hybrid DnCNN (g = 1 nS), and Hybrid DnCNN (g = 5 nS), respectively. Error bars indicate standard deviation.

As we expected, our example case drawn from the image set shows that the G-filter smoothes the background noise and contour information, but the DnCNN compensation can repair the contour and further smooth the background (Fig. 5B). In stark comparison, DnCNN alone only keeps image contours but cannot reduce salt-and-pepper noise. Additionally, the blind denoising performance of Hybrid DnCNN outperforms the original DnCNN in both PSNR and SSIM indices, especially on sizeable blind noise (Fig. 5C). For example, under 14 dB original image quality, the denoising performance of Hybrid DnCNN increases by ∼ 20% on PSNR and by ∼ 45% on SSIM than DnCNN alone. Furthermore, decreasing the gap junction plasticity from 5*nS* to 1*nS* slightly improves image quality on both PSNR and SSIM, suggesting the new denoising architecture is insensitive to the choice of gap-junction strength. The steady performance improvement on PSNR (∼ 2% growth rate) and SSIM (∼ 3% growth rate) with the increase of noise intensity reflects the robustness of Hybrid DnCNN.

### The gap junction strength modulates the receptive field of Hybrid DnCNN

How does the G-filter improve DnCNN’s ability in blind denoising? We consider G-filter as the abstraction of the first layer in the retina, and its gap junction conductance is modulated by external environment – light conditions ***Bloomfield and Völgyi (2009); Hu et al. (2010***). Changes in the G-filter’s receptive field (Fig. 1E) reflect its adaptation to the external environment. Such an adaptation will be passed layer by layer, and eventually, the visual cortex (DnCNN) is likely influenced. Therefore, we speculate that the G-filter may impact the receptive fields of neurons in the higher visual hierarchy - the visual cortex.

Next, we attempt to investigate the effects of G-filter on the receptive fields of the Hybrid DnCNN model through a modified attribution map visualization ***Baehrens et al. (2010); Simonyan et al. (2013***). The attribution map identifies the pixels in the original image that contribute to the value of a chosen pixel in the output layer of the Hybrid DnCNN model, which is analogous to the concept of the receptive field in neuroscience. In brief, the attribution map is obtained through “Integrated Gradients”, a path attribution method ***Sundararajan et al. (2017***). Initially, it was designed for classification tasks requiring “prediction neurons”. To make Integrated Gradients applicable to the Hybrid DnCNN model, we set each neuron in the output layer as a graded prediction neuron. For an *m* × *n* input image, the number of output labels is equal to the number of pixels in the input image (mn) (see “Methods”) (Fig. 6A), while each output neuron that encodes a single pixel in the output image has an attribution map. The results show that the size of the attribution map of an output neuron in the Hybrid DnCNN increases with gap junction strength, similar to how the STA receptive field of a correspondent photoreceptor varies with the strength of gap junctions (Fig. 6B, C).

**Figure 6.**
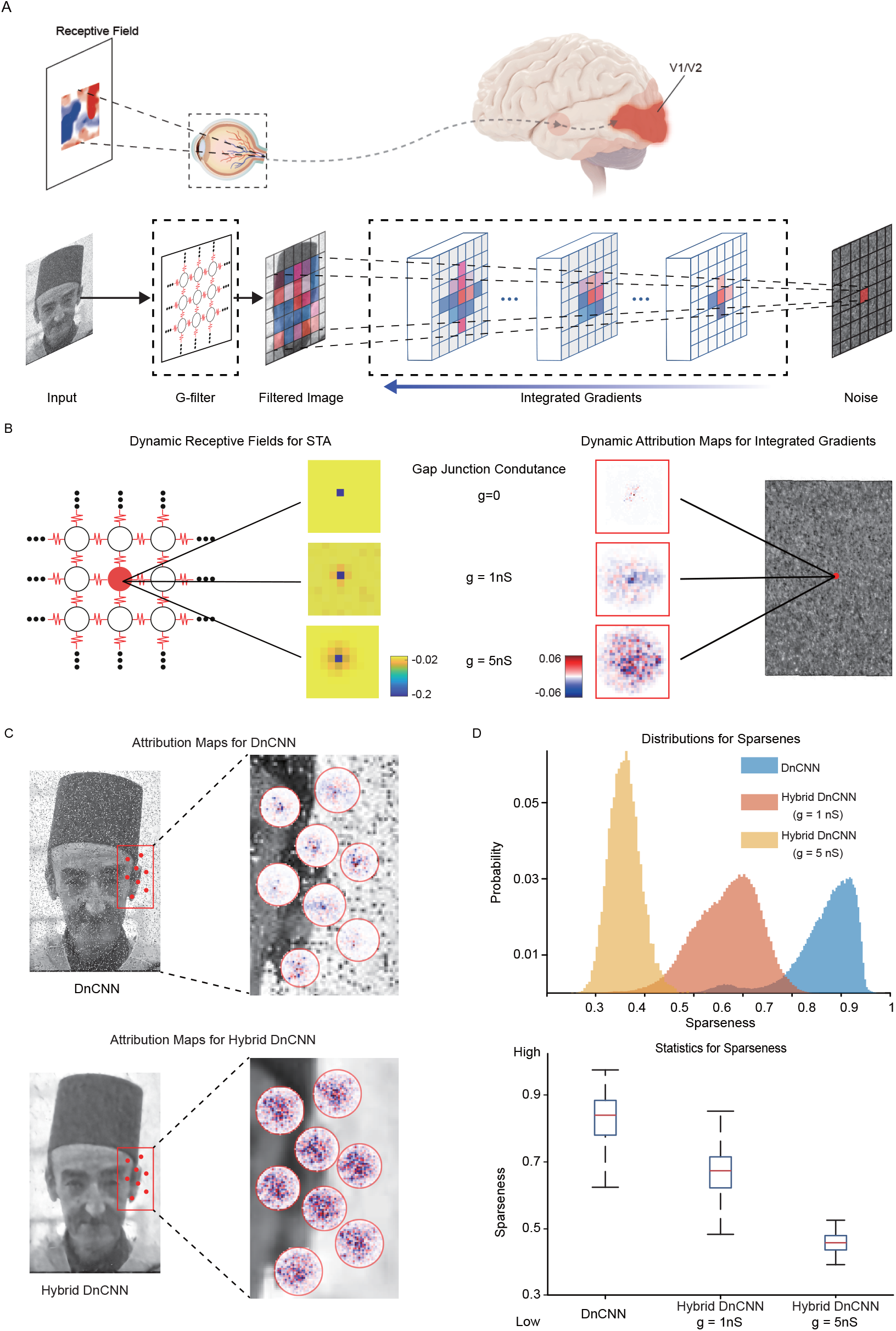
The attribution maps (receptive fields) of the Hybrid DnCNN show consistency with the receptive fields of the photoreceptor gap junction network. (A) The integrated gradients method for evaluating neuron’s receptive field (attribution map) in Hybrid DnCNN. The method is similar to measuring the receptive fields of neurons in physiological experiments. (B) The Comparison of neurons’ receptive fields in the G-filter and the output layer of Hybrid DnCNN. Their receptive field scopes are synchronously modulated by gap junction conductance. (C) Attribution maps of DnCNN and Hybrid DnCNN for a sample image. The red dots indicate pixels whose attribution maps are evaluated by the integrated gradient method. (D) Sparseness statistics of attribution maps of DnCNN and Hybrid DnCNN on all test images. Above is the sparseness distribution of the attribution maps, and below is a boxplot of the sparseness, where the sparseness decreases with the gap junction conductance

Finally, to obtain a quantitative analysis of the attribution map in output neurons, we introduce a measure called “sparseness” to characterize the scope of the attribution map ***Hoyer (2004***) (see “Methods”). Higher sparseness means that fewer pixels in the original image have high contributions (weights) to the output neuron, which makes the attribution map look “sparse”. By using this metric, we find that the attribution maps of neurons in the output layer have high sparseness (0.85) in the absence of the G-filter (the gap junction conductance is 0*nS*) and low sparseness (0.68) in the presence of the G-filter (5*nS*). Noticeably, further increasing the conductance (from 5*nS* to 10*nS*) will make the attribution map denser (from 0.68 to 0.46) (Fig. 6D), but it may not necessarily mean that the denser, the better. Instead, a moderate conductance will achieve the best performance. Thus, by introducing gap junctions and setting a moderate conductance, the sparseness is properly reduced, thereby increasing the number of pixels in the input image that are used as references for blind denoising in the output layer. This enables the Hybrid DnCNN to more accurately obtain the distribution and intensity of noise during learning. In addition, although the receptive fields of pixels in Hybrid DnCNN are more extensive than that of photoreceptors (Fig. 1A), the trends in these receptive fields are consistent; both expanded with increasing gap junction conductance (Fig. 1E and 6D). Considering the attribution map is computed after training the DnCNN, it appears that the model learned to adapt to the plasticity of gap junctions.

### G-filter strengthens DCNNs’ defense against blind noise in classification tasks

Many state-of-the-art (SOTA) DCNNs can easily beat human observers in image classification tasks ***Szegedy et al. (2016); Simonyan and Zisserman (2014); He et al. (2016***). However, the human visual system appears to be more robust than DCNNs when attacked by blind noise. In a psychophysical experiment, the human observers were asked to perform a forced-choice noisy image categorization task and compared the performance of DCNNs in the same task ***Geirhos et al. (2018***). In this experiment, the noise distribution in the training dataset is different from that in the test dataset, which is a typical example of a blind noise attack. The experiment results show the human observers significantly outperform vanilla DCNNs in this specific classification task. Since the G-filter enhances the DCNN’s generalization ability in the previous image-denoising-and-reconstruction task, could the G-filter also improve the DCNN’s ability to resist noise in more general tasks, such as image classifications?

Next, we combine G-filter with the classic DCNNs (Hybrid DCNNs), including ResNet50 ***He et al. (2016***), VGG19 ***Simonyan and Zisserman (2014***) and Inception V3 ***Szegedy et al. (2016***) (Fig. 7A). These DCNNs are very deep and powerful networks, having won the prestigious ImageNet Classification Challenge. We then follow the procedures in the psychophysical experiment. Specifically, we conduct experiments by training Hybrid DCNNs on images with different intensities of salt and pepper noise (noise percentage of 10%, 20%, 35%, 50%, 65%, 80% and 95%) and testing them on images with different levels of uniform noise (noise width with 0.03, 0.05, 0.1, 0.2, 0.35, 0.6 and 0.9) (Fig. 7B, C) (see “Methods”). From previous results ***Geirhos et al. (2018***), the classification accuracies of ResNet50, Inception V3, and VGG19 all significantly drop when increasing Uniform noise width. At the same time, human subjects show a slower downward trend until, in extreme cases, they fell to the same accuracy as DCNNs. (blue lines and dashed black lines in Fig. 7D). Expectedly, the Sal-tand-Pepper-noise-trained Hybrid DCNNs are much more defensive to the Uniform noise, which is the blind noise to Hybrid DCNNs (solid red lines in Fig. 7D). Surprisingly, as the noise width becomes larger (0.6, 0.6, and 0.4 for ResNet50, Inception V3, and VGG19, respectively), the accuracy of Hybrid DCNNs far exceeds that of humans, at which point human observers’ judgments gradually becomes unreliable, ultimately up to guessing (Fig. 7D). Therefore, the combination of G-filter and DCNNs conducts a stronger blind noise defense than classic DCNNs, providing a better solution to problems more likely to be encountered in the real natural environment. Moreover, as these large DCNNs resemble deep layers of the visual hierarchy (Fig. 7A), the results of the hybrid DCNNs suggests that retina gap junctions may also impact the performance of higher visual centers.

**Figure 7.**
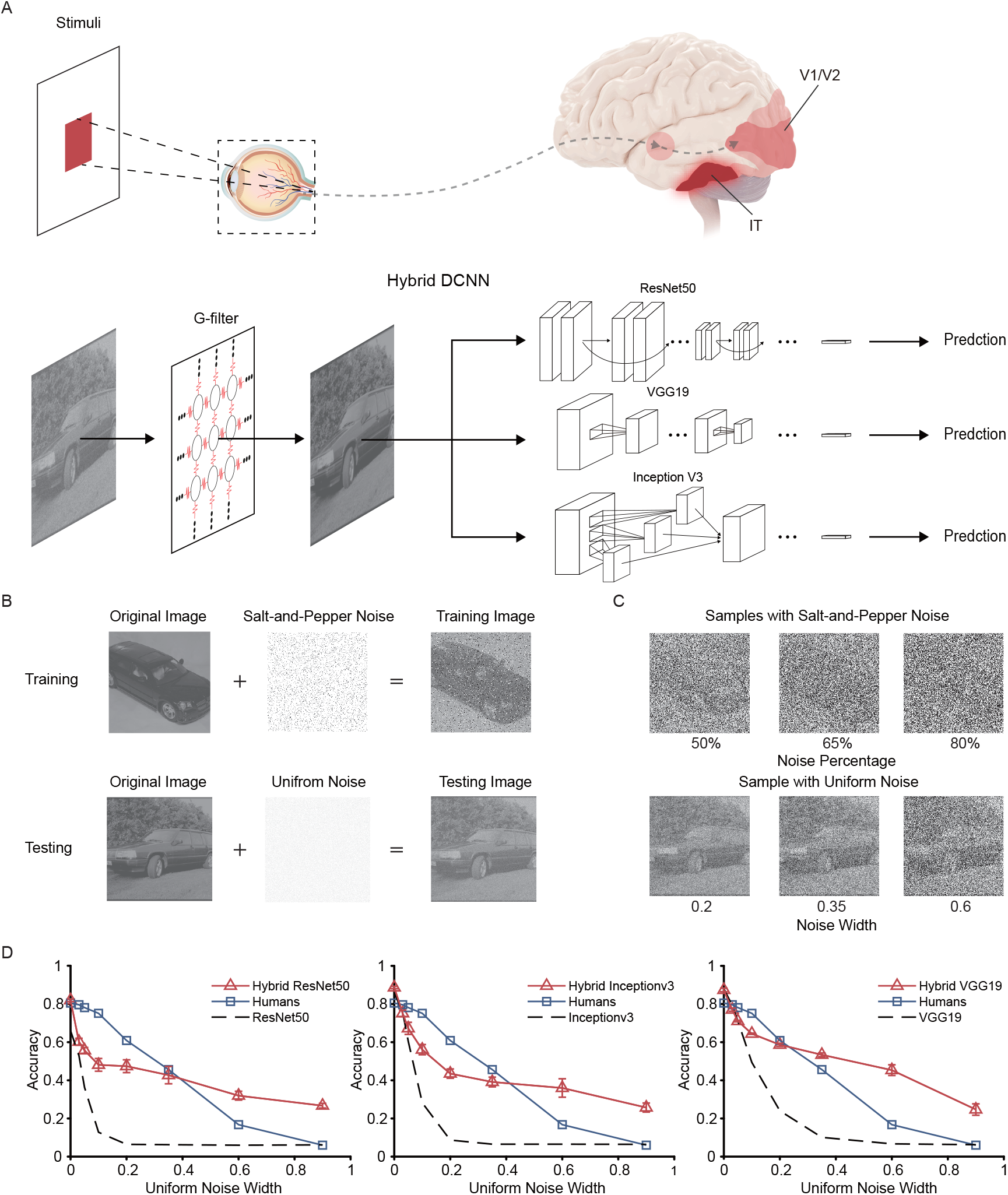
The combinations of G-filter and classic DCNNs (Hybrid DCNNs) simulate the object recognition in the human visual system, and enhances the performance against blind noise. (A) The architectures and workflows of Hybrid DCNNs. The G-filter is a computational module placed in front of ResNet50, VGG19 and Inception V3 for defending blind noise in object recognition task. The G-filter is an abstraction of the retina, while the DCNNs are related to the ventral visual pathway (from V1 to IT) in the cortex. (B) Illustration of training and testing set. The dataset and experiment settings are from Geirhos et al. (2018). This dataset is a subset of the ImageNet dataset with 16 categories. The network learns from a training set with Salt-and-Pepper noise and predicts from a testing set with uniform noise. (C) Several noisy samples for training and testing. The Salt-and-Pepper noise levels (noise percentage) of samples in training set are 50%, 65%, 80%. The Uniform noise levels (noise width) of samples in testing set are 0.2, 0.35, 0.6. (D) The classification accuracies of different models under different Uniform noise levels. Dash lines represent the prediction accuracies of different DCNNs under different noise levels as baseline. Black lines represent human’s prediction accuracies, which comes from a psychological experiment Geirhos et al. (2018). Red lines represent the prediction accuracies of Hybrid DCNNs. Note that Hybrid DCNNs stably surpasses baseline and with the increase of noise levels, the accuracies of Hybrid DCNNs even surpass human.

## Discussion

In this work, we combine the photoreceptor network with several well-established machine learning architectures ***Zhang et al. (2017); Szegedy et al. (2016); Simonyan and Zisserman (2014); He et al. (2016***) and demonstrate its overall improvement. Adjacent neurons in the photoreceptor network can mutually share input information to the extent that varies with the electrical coupling strength of gap junctions. We exploit this property and modify it as a layer of machine learning structure, called gap junction filter (G-filter). G-filter appears the most general denoiser compared with other standard noise filters under multiple noise types. Intriguingly, G-filter can transform noise distributions into a similar bell-shaped form, regardless of the original forms, providing it the ability to resist blind noise. We next integrate G-filter with several DCNNs to slightly mimic the whole visual hierarchy and evaluate the enhancement of this denoising layer on two important visual tasks - image reconstruction and object recognition. In the image reconstruction task, we combine G-filter with a classic convolutional denoising model (DnCNN) and dramatically improve the blind denoising performance. With further investigation, we find the output neurons in DnCNN learn to adapt to the gap junction plasticity. In the object recognition task, our results show that the Hybrid DCNN models reach the performance of human subjects on classification with moderate blind noise interference and even surpass human subjects under large blind noise. Overall, the G-filter can considerably facilitate blind denoising in the visual hierarchy.

### Comparison with denoising methods in computer vision

Denoising is a classical problem in machine learning, which has long been studied over the last decades ***Fan et al. (2019); Gu and Timofte (2019); Goyal et al. (2020***). Denoising methods can be roughly classified into three categories: 1) classic denoising methods, 2) transform domain methods, and 3) deep learning-based methods.

Many classic denoising methods***Lebrun et al. (2012***) aim to remove noise from the spatial domain, eliminating noises in the original image by pixel correlation filters, such as Gaussian, median, and average filters. These methods can remarkably deal with their corresponding noise, such as Gaussian filter versus Gaussian noise and median filter versus Salt-and-Pepper noise. Transform domain methods***Xiong et al. (1999); Dabov et al. (2006); Lebrun et al. (2012); Shao et al. (2013***) initially transform the spatial domain into the frequency domain by the Fourier transform and cut off high-frequency signals to remove noise. Since then, various transform domain methods have been established, such as block-matching and 3D filtering (BM3D)***Dabov et al. (2006***). With the rising popularity of deep learning, several deep learning-based methods ***Zhang et al. (2017); Guo et al. (2019); Tian et al. (2020); Wu and Gao (2021***) have been recently introduced into the image denoising field. One classic deep learning-based denoising method is Denoising Convolutional Neural Network (DnCNN) ***Zhang et al. (2017***), which leverages residual learning to denoise. DnCNN trains the network to learn noise and subtracts noise from the noisy image to acquire the denoised image. Although these methods have achieved remarkable denoising performance, they show weak effectiveness when tackling blind noise because they are overly sensitive to a hyperparameter – noise level estimation *σ*. It causes the denoising methods under given level *σ* to be unsuitable for other noise levels. To adaptively evaluate noise level, a complex additional convolutional subnetwork ***Guo et al. (2019***) has been introduced. However, the computation cost becomes significantly expensive.

In contrast to these denoising methods, the G-filters is designed with strict biological constraints, including the voltage waveform and the gap junction conductance range, where only the gap junction conductance is a tunable hyperparameter. Intriguingly, although tuning of gap junction conductance affects image quality in low-level image processing, this dependency is significantly reduced in high-level image processing when combined with deep learning models. It is worth noting that we did not fine-tune the parameters to achieve high performance, but only test two values (1*nS* and 5*nS*), which shows the high robustness of the biological visual system.

### The denoising performance of the gap junction network follows the circadian rhythm

An unusual phenomenon of the G-filter is that the high contrast level of images will impair its denoising performance, as demonstrated in Fig. 3. In our daily life, high contrast is usually related to good lighting conditions, and changing the lighting conditions from bright to dim will reduce the clarity of the outline of the object, thereby reducing the dependence of the image contour and more reflecting the denoising effect of the G-filter. Since this is an intrinsic feature of gap junction networks, the retina has evolved to accommodate this feature. The circadian change of light intensities will regulate the concentrations of dopamine and nitric oxide (NO) in the retina, resulting in the open of gap junctions in dim light and close in bright light ***Bloomfield and Völgyi (2009***). In contrast, although gap junctions also exist in the cortex, such as between interneurons ***Traub et al. (2001***), the conductance of those gap junctions is mediated by chemical receptors but not the diurnal rhythm of light ***Kawaguchi and Kubota (1997); Turecek et al. (2014***). Therefore, we hypothesize that the denoising performance of G-filter is also in accordance with the change of natural light conditions.

To verify our hypothesis, we consider a day-night dataset ***Zhou et al. (2016***), which incorporates continuous shooting images (48 hours). The dataset is added with large Salt-and-Pepper noise to represent the noisy environment (See “Supplementary” for detail). The gap junction conductance of G-filter is forced open in our experiment and does not change with light conditions. The results show that the denoising performance of G-filter oscillate with the diurnal rhythm and achieves the highest performance at night. This result could be biologically plausible if the gap junctions are kept open in the retina, and perhaps the reason why the retina adjust the conductance of gap junctions through day and night.

### The gap junction network in the retina may facilitate dependencies or correlations among noises

Another important finding of G-filter is that it can transform noise distribution into a similar form, which may significantly facilitate noise reduction in the subsequent networks. Biologically, keeping correlations among noises in the input signals can be beneficial for the signal processing performed by the visual cortex. Several computational and biological researches ***Pillow et al. (2008); Zylberberg et al. (2016); Ruda et al. (2020***) have attempted to address the functional roles of noise correlation. In previous work, the model fitted by the recordings in vitro from ON and OFF parasol ganglion cells shows that statistical dependencies in the responses of sensory neurons can provide additional sensory information ***Pillow et al. (2008***). As a result, decoding under the dependence assumption can extract 20% more information than the independence assumption***Pillow et al. (2008***). Another biophysical experiment on ON-OFF directionally selective retinal ganglion cells shows that noise correlation has a strong stimulus dependence, influencing population coding by co-shaping signals and noise ***Zylberberg et al. (2016***). The computational model based on this experiment present that stimulus-dependent noise correlations significantly promote the precision of motion direction encoding by twice as much as independent noise ***Zylberberg et al. (2016***). Additionally, a GLM-based model fitted by RGC responses of the rat retina segments shows that If the noise is independent across neurons, the brain will suffer a failure of retinal population codes ***Ruda et al. (2020***). Therefore, the ability to transform noises into homogeneous forms illustrated by G-filter will likely promote the correlations of noises in retinal population coding. In this way, the retina can increase the spatial-temporal correlations of noises in population responses and facilitate the visual cortex to filter out these noises.

### The implication for retina disease

The contour repairment effect of the Hybrid DCNNs is correspondent to the visual cortex’s compensatory mechanisms after receiving signals from the retina. The human brain has powerful compensatory mechanisms to recover visual information that are lost with retina injuries ***Baker et al. (2005); Dilks et al. (2014***). However, these mechanisms can also bring some side effects. Due to the brain’s strong adaptability, minor lesions will be disguised, and patients might not have any noticeable signs in the early phase of retina disease, which is inconvenient to early phase diagnoses. For example, A major disease caused by retinal damage is macular degeneration (MD) ***Jager et al. (2008); Lim et al. (2012***). The macula is an area near the retina’s center, responsible for the central, high-acuity, and color vision, containing high density of cones ***Kolb (2012***). Macular degeneration, which is heavily related to age, is an eye disease that can gradually blur central vision ***Lim et al. (2012***) and severely lowers daily life quality and working ability. However, due to the automatic compensatory reorganization of the visual cortex, the early or intermediate MD is usually imperceptible ***Baker et al. (2005***). Concretely, as the visual cortex is retinotopically organized, the loss of central vision in the retina will deprive the input of a specific region in the visual cortex that are supposed to respond to central stimuli in normal conditions ***Baker et al. (2005); Dilks et al. (2009***, 2014). To recover the foveal vision, the visual cortex can adapts to reorganize the functional properties of the deprived cortex ***Baker et al. (2005***).

In our experiments, as indicated by “Integrated Gradients” analysis, the internal representations of deep neural network models also alter with the presence or absence of G-filter. If salt-and-pepper noise is added to the G-filter, such situation can be considered as damaged photoreceptors in the retina. Comparably, the DCNN parts in our hybrid models show similar functional reorganization as in the real visual cortex (Fig. 7). Therefore, the hybrid models have the potentials to serve as retinal disease models in simulation.

### The inspiration for deep learning

Learning from the retina is likely indispensable in designing deep learning frameworks for visual processing. Recently, many image denoising methods developed in deep learning field are inspired by neuroscience ***Carandini and Heeger (2012); Ballé et al. (2015); Huang et al. (2019***). For example, a biological phenomenon called local normalization, which is defined as neurons in the visual cortex suppressing their neighboring neurons, has been successfully applied in image denoising***Carandini and Heeger (2012); Ballé et al. (2015***). Later on, the top-down feedback pathways in the visual cortex inspired a novel DCNN architecture, which can restore noisy images with high reconstruction accuracy ***Huang et al. (2019***). However, current methods usually neglect the importance of the retina and only consider it as a simple filter. Our results show that the photoreceptor gap junction network as one part of the retina, rather than a simple filter, has significant contributions in blind denoising, reshaping noise distribution, and generally improving the performance of various learning tasks. Hence, the introduction of gap junction may provide a new angle, i.e., the importance of the retina, for the future design of machine learning frameworks. Besides, the G-filter we proposed only involves one layer of the retina, nevertheless gap junctions widely exist at all layers. With the introduction of other layers, we can build a deep retina block with gap junctions. The combination of such a retina block with DCNNs may greatly enhance the abilities of noise reduction and feature extraction. Overall, our G-filter is an excellent blind denoiser strictly designed following biological mechanisms and successfully boosts machine learning models’ potentialities acting in the simulated visual hierarchy.

## Methods and Materials

We consider a rod photoreceptor network with grid-like connectivity (Fig. 2B). Photoreceptors are modeled by the Hodgkin-Huxley equations. The simulation platform is NEURON ***Hines and Carnevale (1997***), and the time-step is dt = 0.5 ms. Photoreceptors are connected by gap junctions (electrical synapses), where gap junction conductance is biologically plausible (0*nS* ∼ 10*nS*). Inspired by the photoreceptor network, we design a gap junction filter for blind denoising, and investigate its role and impact on deep neural networks. The model is written in Python and uses the Pytorch library ***Paszke et al. (2019***).

### Photoreceptor Model

We model the photoreceptor with Hodgkin-Huxley equations, which offers high simulation precision. It is adapted from previous works ***Baylor et al. (1984); Baylor and Nunn (1986); Kamiyama et al. (1996); Torre et al. (1990***), and we retune the parameters in the NEURON simulator. This model is considered as a single compartment model, which only has one segment with length *L* = 10*µm* and diameter 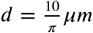. The following ordinary differential equation describes the membrane voltage (*v*_*m*_) of the photoreceptor:

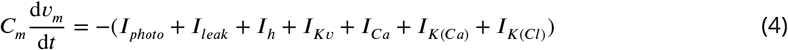

where the initial membrane voltage is *v*^0^ = −41.42868*mV*, the membrane capacitance is *C*_*m*_ = 30*µF* /*cm*^2^, *I*_*photo*_ is the photocurrent, *I*_*leak*_ is the leakage current, *I*_*h*_ is the hyperpolarization-activated current, *I v* is the delayed rectifying potassium current, *I*_*Ca*_ is the calcium current, *I*_(*Ca*)_ is the Cadependent potassium current, and *I*_(*Cl*)_ is the Ca-dependent chloride current. Currents in the photoreceptor model are decomposed into photocurrent (*I*_*photo*_), and ion channel currents (*I*_*leak*_, *I*_*h*_, *I*_*K v*_, *I*_*Ca*_, *I*_*K* (*Ca*)_, and *I*_*K* (*Cl*)_).

The photocurrent (*I*_*photo*_) involves a complex phototransduction cascade as illustrated in Fig.8, following the previous work ***Kamiyama et al. (1996***). In the model of phototransduction, input photons trigger a series of biophysical processes and finally influence the concentration of cyclic guanosine monophosphate (*cGMP*), which controls ionic flow (their parameters are reported in Table1).

**Table 1.** The parameter of phototransduction mechanism

| Parameter | Value |
| --- | --- |
| $\alpha_1$ | $0.05 \text{ ms}^{-1}$ |
| $\alpha_2$ | $0.0000003 \text{ ms}^{-1}$ |
| $\alpha_3$ | $0.00003 \text{ ms}^{-1}$ |
| $\varepsilon$ | $0.0005 \text{ uM}^{-1} \text{ ms}^{-1}$ |
| $T_{Tot}$ | $1000 \text{ uM}$ |
| $\beta$ | $0.00025 \text{ uM ms}^{-1} \text{ pA}^{-1}$ |
| $\tau_1$ | $0.0002 \text{ uM}^{-1} \text{ ms}^{-1}$ |
| $\tau_2$ | $0.005 \text{ ms}^{-1}$ |
| $PDE_{Tot}$ | $100 \text{ uM}$ |
| $\sigma$ | $0.001 \text{ uM}^{-1} \text{ ms}^{-1}$ |
| $A_{max}$ | $0.0656 \text{ uM ms}^{-1}$ |
| $K_c$ | $0.1 \text{ uM}$ |
| $V$ | $0.0004 \text{ ms}^{-1}$ |
| $k_1$ | $0.0002 \text{ uM}^{-1} \text{ ms}^{-1}$ |
| $k_2$ | $0.0008 \text{ ms}^{-1}$ |
| $e_r$ | $500 \text{ uM}$ |
| $b$ | $0.00025 \text{ uM ms}^{-1} \text{ pA}^{-1}$ |
| $\gamma_{Ca}$ | $0.05 \text{ ms}^{-1}$ |
| $C_{a0}$ | $0.1 \text{ uM}$ |
| $J_{max}$ | $5040 \text{ pA}$ |
| $K$ | $10 \text{ uM}$ |
| Initial parameter | Value |
| $Ca_0$ | $0.3 \text{ uM}$ |
| $Ca_{b0}$ | $34.9 \text{ uM}$ |
| $cGMP_0^*$ | $2 \text{ uM ms}^{-1}$ |

**Figure 8.**
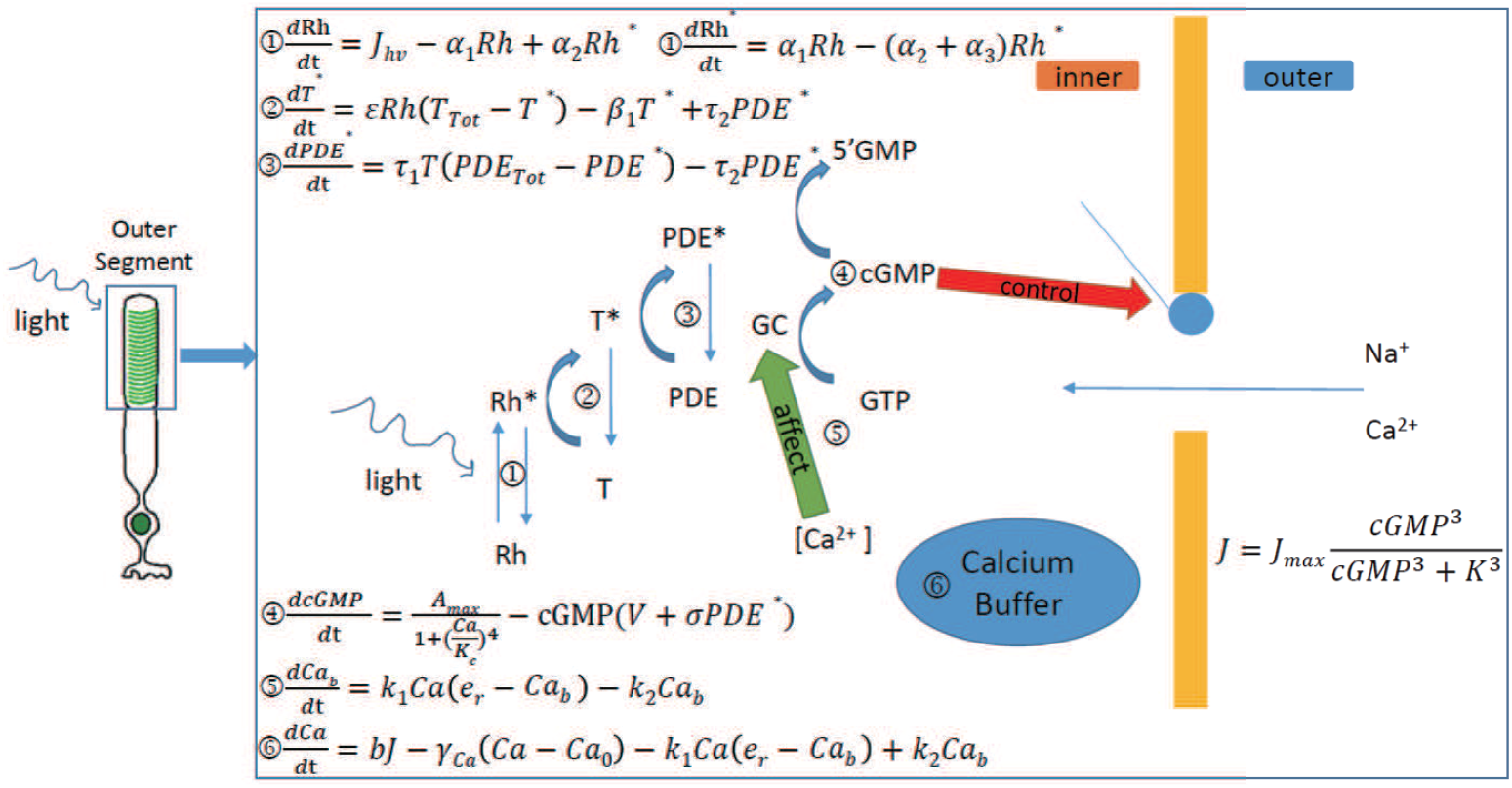
The mechanism of phototransduction

The cyclic GMP-activated ionic current (*J*) is a function of the concentration of cyclic GMP (*cGMP*) and the photocurrent (*I*_*photo*_) is proportional to cyclic GMP-activated current. Therefore, *I*_*photo*_ is expressed as:

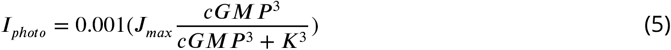

Except for the photocurrent (*I*_*photo*_), other ion channel currents *I*_*ion*_ (*I*_*leak*_, *I*_*h*_, *I*_*Kv*_, *I*_*Ca*_, *I*_*K* (*Ca*)_, and *I*_*K*(C*l*)_) can be described by Hodgkin-Huxley equations of the form:

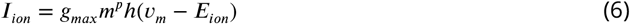

where *g*_*max*_ is the maximum conductance, *m* is the activation variable, *h* is the inactivation variable, *p* is the gating exponent and *E*_*ion*_ is the reversal potential. These ion channels are summarized in Table2.

**Table 2.** The ion mechanism of photoreceptor

| Mechanism | Conductance Value |
| --- | --- |
| Calcium channel | $2 \text{ mS/cm}^2$ |
| Ca-dependent potassium Channel | $0.5 \text{ mS/cm}^2$ |
| Delayed rectifying potassium Channel | $2.0 \text{ mS/cm}^2$ |
| Noninactivating potassium Channel | $0.85 \text{ mS/cm}^2$ |
| Nonselective cation Channel | $3.5 \text{ mS/cm}^2$ |
| Leak Channel | $0.6 \text{ mS/cm}^2$ |
| Calcium Buffer | - |

Because of the Calcium-dependent currents, we introduce a Calcium buffer (intracellular Calcium concentration system), following previous work ***Kamiyama et al. (1996***). It is available on ModelDB (https://senselab.med.yale.edu/ModelDB/ShowModel?model=95870).

After introducing phototransduction cascade, ion channels and Calcium buffer, the rod photoreceptor model can simulate light responses of photocurrent, which approximates experimental data ***Baylor and Nunn (1986***) (Fig.9).

**Figure 9.**
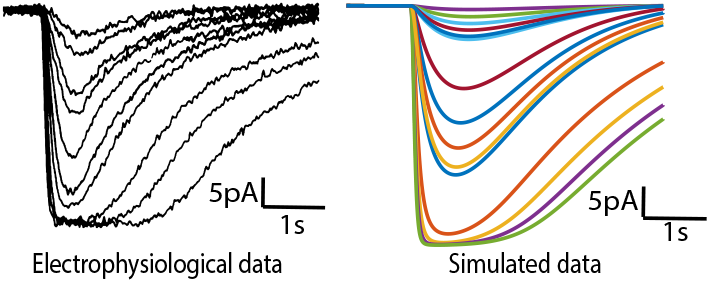
Photocurrents in physiological data (left, adapted from***Baylor and Nunn (1986***)) and our detailed model (right).

### Photoreceptor Network

We consider a one-layer biologically detailed network model, where gap junctions connect the neighboring rod photoreceptors (Fig.2B). The whole network is a 10 × 10 grid structure, where each photoreceptor locates in the intersection point. Thus, the inner current 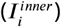 of each photoreceptor *i* can be decomposed into seven components:

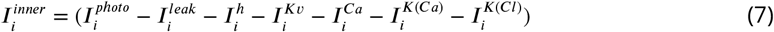

The photoreceptor *i* also receives the gap junction current 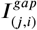 from its neighboring photoreceptor j, which can be described as:

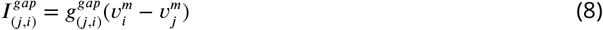

where 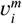 is the membrane voltage of photoreceptor 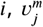 is the membrane voltage of photoreceptor 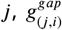 is the conductance of gap junction between photoreceptor *i* and photoreceptor *j*. Therefore, the membrane voltage of each photoreceptor in the network follows

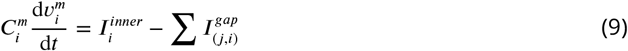

where the initial membrane voltage is 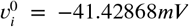, the membrane capacitance is 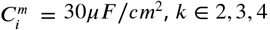.

### STA Analysis

The spike-triggered average (STA) provides an estimate of a neuron’s linear receptive field through the relationship between stimulus and response, which requires stimuli to be strictly “spherically symmetric”, such as Gaussian white noise ***Sharpee et al. (2004***). Because the photoreceptor is a non-spike neuron, we consider each negative peak event (the red point) in voltage response as a ‘spike’ (Fig.10).

**Figure 10.**
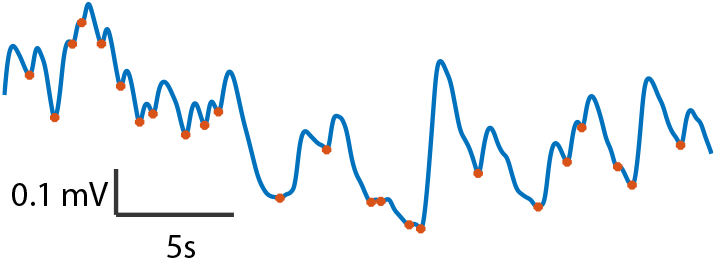
Sample voltage response trace under white Gaussian noise. Red marker donates the peak point (viewed as ‘spike’) in trace.

To compute STA, each photoreceptor in the network is stimulated with white Gaussian noise at 5*Hz*. The whole stimulus time is divided into *k* bins. Let _*i*_ denotes the stimulus vector preceding *ith* bin, and *y*_*i*_ denotes the number of events in *ith* bin. The STA is given by

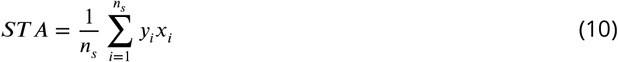

where *n*_s_ is the total number of spikes.

### Gap Junction Filter

Inspired by the information integration function of the rod photoreceptor network, we propose the gap junction filter (G-filter), which can be extended to *m* × *n* grid structure for blind denoising. Each neuron in G-filter is a single compartment model, which has one segment with length *L* = 10*µm* and diameter 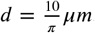. The currents of each neuron in the G-filter consist of two parts: the inner current and the gap junction current. The inner current 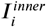 of each neuron *i* in the G-filter follows:

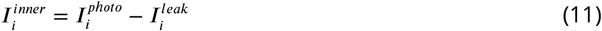

where 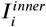 is defined as in Eq.3. The gap junction current 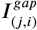 between neuron *i* and neuron *j* is the same as in the photoreceptor network. Therefore, the membrane voltage 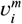 of each neuron *I* in the G-filter follows

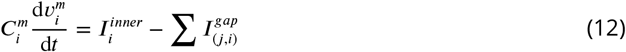

where the initial membrane voltage is 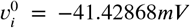 the membrane capacitance is 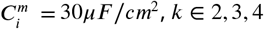.

### The Image Processing of Gap Junction Filter

For the *m* × *n* input image *M*, the G-filter has the same neurons with *M*. Each pixel *M*_*ij*_ ∈ [0, 1] is assigned to *g*_*max*_ in Eq.3, and generate a current curve *I*_*ijt*_, where the delta time is 1*m* and the duration time *t* is 2. Then, the current is injected into corresponding neuron in the G-filter and the membrane voltage *V*_*ijt*_ is recorded. All membrane voltages *V* during time *t* is a *m* × *n* × *t* matrix. We first get the maximum voltages *V*_*max*_ by time *t*. Then we normalize *V*_*max*_ to [0, 1] and reconstruct the filtered image.

### Other Filters

The Gaussian Filter first generate a 3 × 3 Gaussian kernel following 2D Gaussian function:

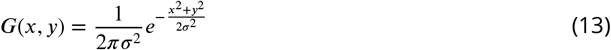

where the sum of all elements in the kernel is 1, and the parameter σ = 0.5. Then it convolutes the kernel with the input image (padding).

The Average Filter first generate a 3 × 3 kernel, where all elements in the kernel are identical and their sum is 1. Then it convolutes the kernel with the input image (padding).

The Median Filter replace each pixel value in the image by the median value of the neighboring pixel, where the window size is 3 × 3.

The Adaptive Median Filter ***Hwang and Haddad (1995***) is an advanced version of the standard one, and its window size is variable. For an image normalized to [0, 1], the selection method of window size follows a two-step process. Firstly, it finds the median value for the current size. Secondly, it checks whether the median value is greater than zero and less than one. If so, the window size is proper or else it changes the window size.

The Max Filter replace each pixel value in the image by the max value of the neighboring pixel, where the window size is 3 × 3.

The Min Filter replace each pixel value in the image by the min value of the neighboring pixel, where the window size is 3 × 3.

### PSNR

The peak signal-to-noise ratio (PSNR) is an expression for the ratio between the signal’s maximum power and the power of noise ***Hore and Ziou (2010***). It is commonly used to evaluate the image reconstruction quality, usually expressed as logarithmic quantity with the decibel (*dB*). The PSNR is given by:

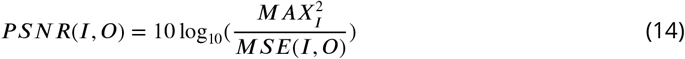

where *I* is the original image, *o* is the noisy image, *MAX*_*I*_ is the maximum possible pixel value of the image *I*. The mean squared error (MSE) is defined as:

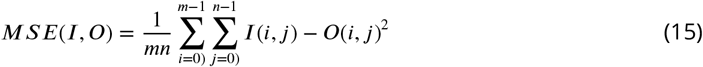

where *m* × *n* is the size of *I* and *o*.

### SSIM

The Structural Similarity Index (SSIM) is a perception-based measure for image reconstruction quality evaluation ***Wang et al. (2004***). The SSIM can be decomposed into three comparison measures between the original image *I* and the noisy image *O*: luminance (*l*(*I, O*)), contrast (*c*(*I, O*)), and structure ((*I, O*)). Thus, the SSIM is defined as:

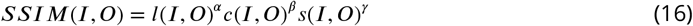

where α, β, and γ are the weights, and the default value is 1. These three measures follow:

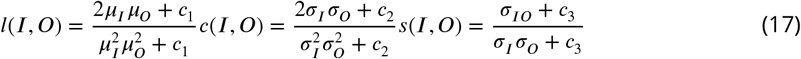

where *µ*_*I*_ is the mean of the original image *I, µ*_*o*_ is the mean of the noisy image 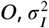 is the variance of the original image 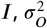 is the variance of the noisy image *o, σ*_*IO*_ is the covariance of the original image *I* and the noisy image *o, c*_1_ = (*k*_1_*L*)^2^, *c*_2_ = (*k*_2_*L*)^2^, and 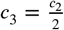 are used to stabilize the division with weak denominator, *L* is the range of the pixel value, *k*_1_ = 0.01, and *k*_2_ = 0.03.

### Noise Generation

The noise types used for testing filter denoising performance include Gaussian noise, Laplacian noise, Salt-and-Pepper noise, Uniform noise, and mixed noise. For each original image *I*, the noise generator *f* is imposed, mapping from *R*^*n*^ to *R*^*n*^. Thus, the noisy image *o* follows:

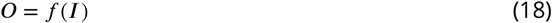

The generating processes of these noises are listed below: For Gaussian noise, the probability density function *f* is given by:

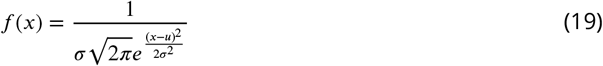

where is the Gaussian random variable, *µ* is the mean value, *σ* is the standard deviation. In this paper, *µ* = 0 and the magnitude of Cf controls the average noise level l of noisy images, where *l* ∈9*d*B, 12*d*B, 15*d*B, 18*d*B.

For Laplacian noise, the probability density function f is given by:

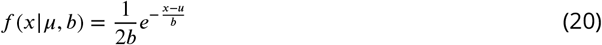

where is the Laplacian random variable, *µ* is the location parameter, *b* is the scale parameter, and *b >* 0. In this paper, the magnitude of *µ* and *b* controls the average noise level *l* of noisy images, where *l* ∈ 9*dB*, 12*dB*, 15*dB*, 18*dB*.

For Salt-and-Pepper noise, each pixel in original image *I* is corrupted with the probability. Half of these corrupted pixels are randomly set to black, and another is randomly set to white. In this paper, the magnitude of r controls the average noise level *l* of noisy images, where *l*∈9*dB*, 12*dB*, 15*dB*, 18*dB*.

For Uniform noise, the probability density function is given by:

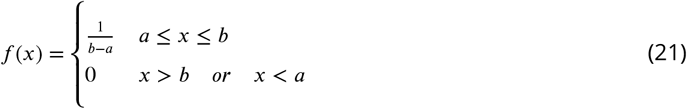

where *x* the uniform random variable, *a* and *b* are the two boundaries. In this paper, the magnitude of *b* − *a* controls the noise level *l* of noisy images, where *l* ∈9*dB*, 12*dB*, 15*dB*, 18*dB*.

The mixed noise is generated by the combination of Gaussian noise, Laplacian noise, Salt-andPepper noise, and uniform noise, which satisfies the average noise level *l* of noisy images, where *l*∈9*dB*, 12*dB*, 15*dB*, 18*dB*.

### The Combination of G-filter and DnCNN for Blind Denoising

In the combination model, noisy images are filtered by G-filter first before being used as training data for DnCNN. We follow DnCNN’s prescription ***Zhang et al. (2017***) in our experiments. We use 400 images of size 180 180 for training. 3, 000 patches of size 50 50 are cropped from the 400 images to train our model. For testing, we use 68 natural images from BSD68. The dataset and code of DnCNN are available on their website (https://github.com/cszn/DnCNN). We train our network using Adam optimizer with an initial learning rate of 0.001. Our network is trained for 50 epochs with a mini-batch size of 128. The learning rate is divided by 10 upon reaching 30 epochs.

### Integrated Gradients Method

The Integral Gradients is the gradient-based attribution method widely applied to explainable artificial intelligence. The Integral Gradients aims to explain the relationship between input and predictions of deep neural networks based on gradients. The output of Integral Gradients is the attribution map, which reflects the contribution of input to the final prediction. The attribution map has the same size as the input image, and the value is between −1 and 1. The positive value in the attribution map shows that the corresponding pixel in the input image has a positive contribution to the final prediction and vice versa. In this paper, the output of DnCNN is not predictions but noise with the size of *mn*. Therefore, we consider each pixel in the output as a prediction. Then, the Hybrid DnCNN has *mn* predictions and we apply the Integral Gradients following previous work ***Sundararajan et al. (2017***).

### Sparseness

The sparseness in this paper refers to the average activation degree of input neurons for each output neuron. Various sparseness measures have been proposed during the last decades. They are mappings from *R*^*n*^ to *R*, where the sparseness is one, if and only if only a single component is non-zero; the sparseness is zero, if and only if all components are equal. In this paper, we use a sparseness measure following previous work ***Hoyer (2004***) to evaluate the attribution map of the output neuron in the combination model of G-filter and DnCNN, which can be described as:

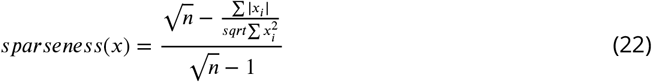

where is the attribution map from Integrated Gradients, *n* is the dimensionality of *x*.

### The Combination of G-filter and DCNNs for Noisy Object Recognition

We use ImageNet (ILSVRC) ***Deng et al. (2009***), which is a large-scale dataset for object classification and detection, for training our Hybrid DCNN models. In order to comparing with human observers ***Geirhos et al. (2018***), we follow their prescription in our experiments, which use the WordNet hierarchy to map ImageNet categories into 16 entry-level categories (airplane, bicycle, boat, car, chair, dog, keyboard, oven, bear, bird, bottle, cat, clock, elephant, knife, truck). In our experiments, the models we used are ResNet50, VGG19, and Inception-V3. Before images are feed into these three networks, G-filter first processes them as shown in Fig.7A.

As the grayscale dataset is used, we add a convolution layer (input channels=1, output channels=3, kernel size=1) to the top of all three neural networks, only for channel expanding. For training, we use SGD with a mini-batch size of 400. The learning rate starts from 0.1 and is divided by 10 when the error plateaus, and the models are trained for 90 iterations. We use a weight decay of 0.0001 and a momentum of 0.9. The training profile is the same in all three networks.

In resnet50, we use basic residual blocks which contain two convolution layers, two batch normalization layer and a ReLU layer. The whole network is constructed by an input-normalize layer, a CONV-BN-RELU module, 3 residual block groups with planes 16, 32, 64, an avgpooling layer and a 16-layer classifier. Our implementation for ResNet50 follows https://github.com/pytorch/vision/blob/main/torchvision/models/resnet.py.

In VGG19, 16 convolutional layers with 3 3 convolution filters is used. The number of channels starts from 64 in the first layer and then increasing by a factor of 2 after each max-pooling layer, until it reaches 512. The network is then followed by three fully-connected layers: the first two have 512 channels each, the third contains 16 channels to performs 16-way classification. The implement is the same as https://github.com/pytorch/vision/blob/main/torchvision/models/vgg.py.

In inception V3, standard inception modules are used. After the feature extracting with a 2048channel linear layer, the network ends with a 16-classifier. We use the network from https://github.com/pytorch/vision/blob/main/torchvision/models/inception.py.

## Appendix 1

### Filters’ Denoising Performance

**Appendix 1 Table 1.** All Filters’ Performance on PSNR

| Noise Type | noisy | Adaptive Median | Average | Gaussian | Max | Median | Min | G-filter |
| --- | --- | --- | --- | --- | --- | --- | --- | --- |
| Gaussian | 9.3931 | 12.3257 | 16.4323 | 16.7648 | 5.6420 | 14.7134 | 7.4840 | **19.4910** |
|  | 12.0365 | 14.9079 | 19.1847 | 19.6653 | 7.6740 | 18.1044 | 8.9699 | **20.9816** |
|  | 14.9113 | 17.6838 | 21.5659 | **22.2076** | 10.2093 | 20.9178 | 10.9068 | 21.7804 |
|  | 18.1941 | 20.7064 | 23.6390 | **24.6108** | 12.9497 | 23.4640 | 13.2402 | 23.3808 |
| Laplacian | 9.5985 | 13.2473 | 16.6411 | 16.9802 | 5.5888 | 15.8758 | 7.4579 | **19.4907** |
|  | 12.0132 | 15.7000 | 19.1456 | 19.6166 | 7.2323 | 19.3830 | 8.7341 | **20.8815** |
|  | 14.4932 | 18.2277 | 21.2598 | 21.8724 | 9.2729 | **21.9666** | 10.3296 | 21.7843 |
|  | 17.8840 | 21.4520 | 23.4909 | 24.4264 | 12.1308 | **24.4911** | 12.6902 | 23.2115 |
| Salt&Pepper | 9.6453 | **28.3222** | 16.5263 | 16.8384 | 5.2340 | 19.8959 | 7.3248 | 18.9588 |
|  | 12.0806 | **30.7300** | 19.0877 | 19.5262 | 6.4557 | 25.8218 | 8.5177 | 20.2686 |
|  | 15.0880 | **32.2882** | 21.6591 | 22.2638 | 8.4892 | 28.2621 | 10.4893 | 21.2754 |
|  | 18.0933 | **32.8780** | 23.6104 | 24.5444 | 10.7724 | 28.8849 | 12.6374 | 22.8880 |
| Uniform | 8.8903 | 11.1816 | 15.8579 | 16.1635 | 5.6221 | 13.1821 | 7.4633 | **19.1190** |
|  | 11.7585 | 13.7830 | 18.9400 | 19.4126 | 8.1607 | 16.5059 | 9.1614 | **20.8624** |
|  | 14.3920 | 16.2600 | 21.1588 | **21.7747** | 10.5340 | 19.0915 | 11.0152 | 21.6886 |
|  | 17.6805 | 19.3204 | 23.3418 | **24.2517** | 13.2001 | 21.9098 | 13.3752 | 23.0765 |
| Blind | 9.5785 | 13.9324 | 16.3319 | 16.6370 | 5.4844 | 16.1899 | 7.4002 | **19.4510** |
|  | 12.4169 | 17.3056 | 19.3707 | 19.8453 | 7.2606 | 20.2933 | 8.8709 | **21.0081** |
|  | 15.2791 | 20.3255 | 21.7421 | 22.3774 | 9.4325 | **23.0744** | 10.8026 | 21.7454 |

**Appendix 1 Table 2.**
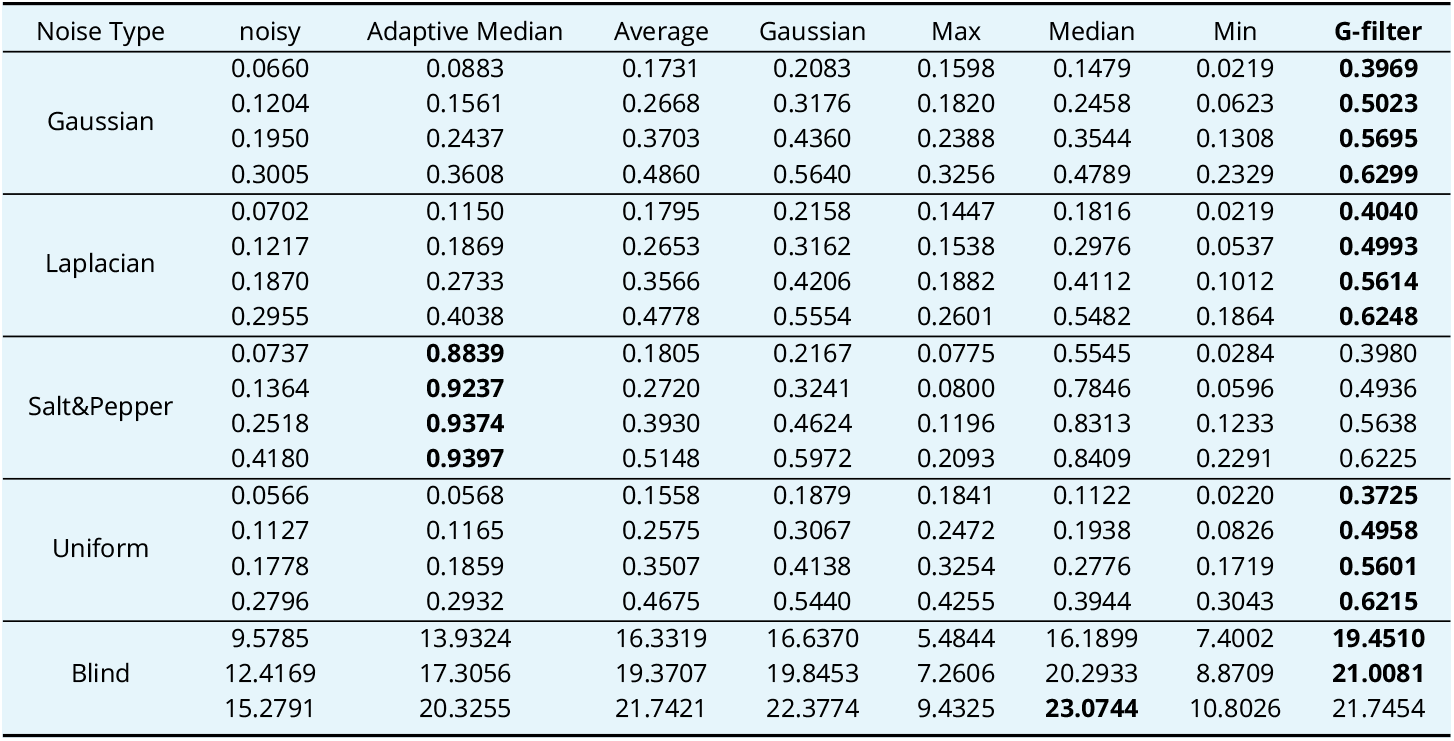
All Filters’ Performance on SSIM

### G-filter and biological circadian rhythm

**Appendix 1 Figure 1.**
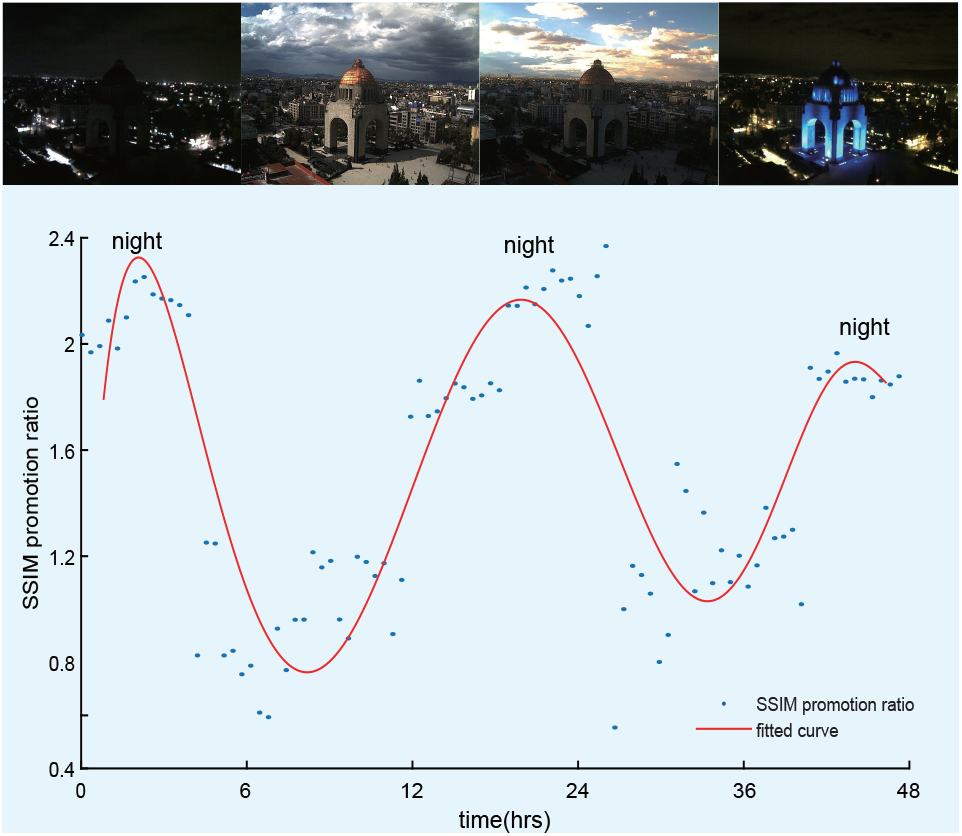
The change of G-filter’s denoising performance is consistent with biological circadian rhythm

We consider a day-night dataset in ***Zhou et al. (2016***), which incorporates continuous shooting images for two days and three nights (48 hrs). The time interval is 15 min. 12 dB’ Salt-and-Pepper noise is added on such a dataset. The results (Fig.8B) show that the denoising performance of G-filter (the conductance is 5 nS) alters with diurnal variation and achieves the highest performance at night (at 375 min, 1615 min, and 2725 min) while decreasing by day (at 935 min and 2250 min). The performance variation of the G-filter is consistent with the biological characteristics of gap junction plasticity. In the daytime, the textures of images are distinct. If we open gap junctions, the image details are smoothed. However, the performance promotion by blind denoising cannot compensate for the information loss by blurring details. Therefore, G-filter’s fixed gap junction conductance will show performance variation consistent with biological circadian rhythm.

### More Samples of Attribution map for DnCNN and Hybrid DnCNN

**Appendix 1 Figure 2.**
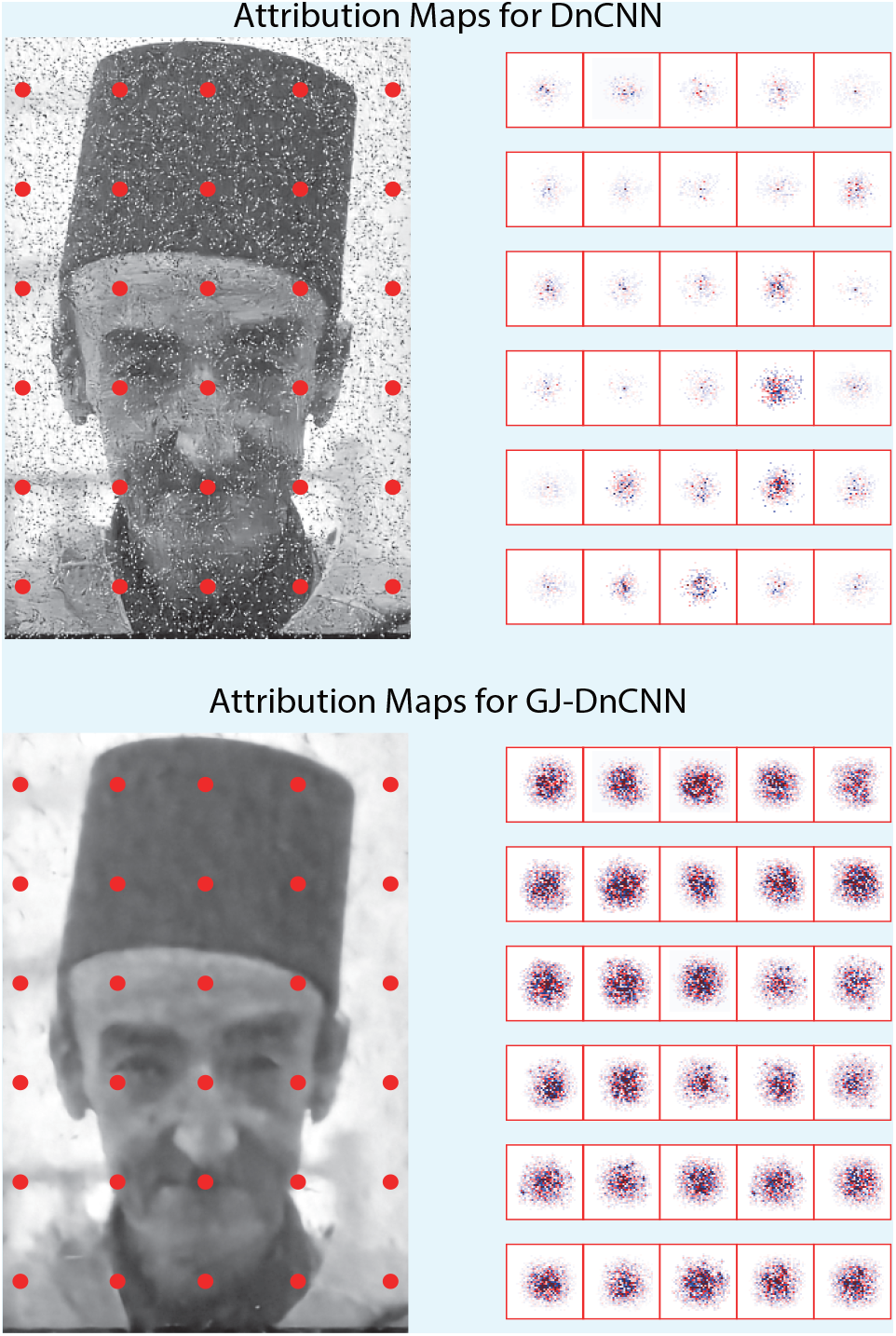
More sample attribution maps of DnCNN without or with G-filter. The red dots indicate the pixels for evaluating their attribution maps by integrated gradients method, and the right is the corresponding attribution maps.

## Notes

### Competing Interest Statement

The authors have declared no competing interest.

